# Genome sequencing and assembly of *Tinospora cordifolia* (Giloy) plant

**DOI:** 10.1101/2021.08.02.454741

**Authors:** Shruti Mahajan, Abhisek Chakraborty, Titas Sil, Vineet K Sharma

## Abstract

During the ongoing COVID-19 pandemic *Tinospora cordifolia* also known as Giloy gained immense popularity and use due to its immunity-boosting function and anti-viral properties. *T. cordifolia* is among the most important medicinal plants that has numerous therapeutic applications in health due to the production of a diverse array of secondary metabolites. Therefore, to gain genomic insights into the medicinal properties of *T. cordifolia*, the first genome sequencing was carried out using 10x Genomics linked read technology and the draft genome assembly comprised of 1.01 Gbp. This is also the first genome sequenced from the plant family Menispermaceae. We also performed the first genome size estimation for *T. cordifolia* and was found to be 1.13 Gbp. The deep sequencing of transcriptome from the leaf tissue was also performed followed by transcriptomic analysis to gain insights into the gene expression and functions. The genome and transcriptome assemblies were used to construct the gene set in *T. cordifolia* that resulted in 19,474 coding gene sequences. Further, the phylogenetic position of *T. cordifolia* was also determined through the construction of a genome-wide phylogenetic tree using 35 other dicot species and one monocot species as an outgroup species.

## INTRODUCTION

*Tinospora cordifolia* is a climbing shrub belonging to the Menispermaceae family that includes more than 400 plant species of high therapeutic properties [1, 2]. It perhaps originated in Africa in the Oligocene epoch (28.57 million years ago) and was spread to Asia in the early Miocene epoch (21.54 million years ago) [3]. *T. cordifolia* is found in tropical and sub-tropical regions including India, China, Sri Lanka, Bangladesh, Myanmar, Thailand, Malaysia, etc. and also known as ‘Giloy’, ‘Amrita’, ‘Guduchi’, and ‘heart leaved moonseed’ [2]. It is a perennial deciduous dioecious plant with morphological characteristics of twining branches, succulent stem with papery bark, alternatively arranged heart-shaped leaves, aerial roots and greenish yellow tiny flowers in the form of racemes inflorescence [2, 4]. Being a climber, *T. cordifolia* needs a supportive plant like *Jatropha curcas* (Jatropha), *Azadirachta indica* (Neem), *Moringa oleifera* (Moringa), etc. for its growth [4]. These co-occurring plants also play an important role in enhancing the production of various secondary metabolites of *T. cordifolia* [4, 5]. Previous reports also indicated the presence of endophytic fungi in the leaves and the stem of this plant but their ecological significance has yet to be studied [6, 7]. This plant produces the secondary metabolites in response to the stress conditions and their concentration also varies based on seasons and its dioecy status [8]. High genetic diversity has been reported in *T. cordifolia* due to the dioecious nature [9-11].

The chemical constituents of this plant have been broadly categorized as alkaloids (tinosporine, magnoflorine, berberine, etc.), terpenoids (tinosporide, furanolactone diterpene, cordifolioside, etc.), phenolics (lignans, flavonoids, phenylpropanoids etc.), polysaccharides (glucose, xylose, rhamnose, etc.), steroids (giloinsterol, ß-sitosterol, etc.), essential oils and aliphatic compounds along with a few other compounds such as giloin, tinosporidine, sinapic acid, tinosporone, tinosporic acid, etc. that are obtained from various parts of the plant [12, 13]. A terpene tinosporaside and an alkaloid berberine were found to be the most dominant compounds in *T. cordifolia* and suggested to use them as its chemical biomarkers [14, 15]. The bitter taste of *T. cordifolia* is due to the presence of tinosporic acid, tinosporol, giloin, giloinin, tinosporide, cordifolide, tinosporin and a few other compounds [12]. A study reported that among the two species of *Tinospora (i*.*e. T. cordifolia* and *T. sinensis*), *T. cordifolia* produces three times higher concentration of berberine than *T. sinensis*, and thus the former is preferred in therapeutics [16].

The bioactive compounds found in *T. cordifolia* have known biological properties such as anti-pyretic, anti-diabetic, anti-inflammatory, anti-microbial, anti-allergic, anti-oxidant, anti-diabetic, anti-toxic, anti-arthritis, anti-osteoporotic, anti-HIV, anti-cancer, hepatoprotective, anti-malarial, and also in immunomodulation etc. [17, 18]. These properties make this species useful in the traditional treatment of several ailments including fevers, cough, diabetes, general debility, ear pains, jaundice, asthma, heart diseases, burning sensation, bone fracture, urinary problems, chronic diarrhoea, dysentery, leucorrhoea, skin diseases, cancer, helminthiasis, leprosy and rheumatoid arthritis. Further pre-clinical and clinical studies have been carried out to indicate its potential to treat leucopenia induced by breast cancer chemotherapy, hepatic disorders, post-menopausal syndrome, obstructive jaundice, etc. [19, 20]. These diverse and important therapeutic applications make it a species of interest for a broad scientific community. Interestingly, this Giloy plant has gained tremendous therapeutic interest and significance during the recent and ongoing COVID-19 pandemic [21].

However, despite the widely known and important medicinal properties of this plant its genome assembly is yet unavailable. A preliminary study reported the transcriptome (482 Mbp data) of this species from leaf and stem tissues using 454 GS-FLX pyrosequencing [22]. A recent karyological study reported 2n=26 as the chromosome number in *T. cordifolia*, which was also supported by the earlier studies [23-25]. Thus, to uncover the genomic basis of its medicinal properties and for further exploration of its therapeutic potential, we carried out the first genome sequencing and assembly of *T. cordifolia* using 10x Genomics linked reads. This is the first draft genome assembly of *T. cordifolia* which is also the first genome sequenced so far from the medicinally important genus *Tinospora* and its family [26]. We also carried out a comprehensive deep sequencing and assembly of the leaf transcriptome using Illumina reads. The genome-wide phylogenetic analysis was also carried out for *T. cordifolia* with other dicot species and a monocot species as an outgroup to determine its phylogenetic position.

## METHODS

### Sample collection, species identification, nucleic acids extraction and sequencing

The plant was brought from a nursery in Bhopal, Madhya Pradesh, India (23.2599° N, 77.4126° E). DNA and RNA extraction were carried out using a cleaned leaf, which was homogenized in liquid nitrogen. The TRIzol reagent was used for RNA extraction [27]. For genomic DNA extraction, the powdered leaf was washed with 70% ethanol and distilled water in order to eliminate any such compounds that may hinder the extraction process and employed CTAB based Carlson lysis buffer for the isolation [28]. Two genes: one nuclear gene and one chloroplast gene (Internal Transcribed Spacer and Maturase K, respectively) were used for the species identification. These genes were amplified and sequenced at in-house sanger sequencing facility. The TruSeq Stranded Total RNA library preparation kit with Ribo-Zero Plant workflow (Illumina, Inc., USA) was deployed for preparing the transcriptomic library. The genomic library for linked reads was prepared using Gel Bead kit and Chromium Genome library kit on a Chromium Controller instrument (10x Genomics, CA, USA). The quality of both the libraries (transcriptomic and genomic) was checked on Tapestation 4150 (Agilent, Santa Clara, CA) and sequenced on an Illumina platform, NovaSeq 6000 (Illumina, Inc., USA) for producing paired end reads. The comprehensive DNA and RNA extraction procedure is mentioned in **Supplementary Text S1**.

### Genome assembly

An array of Python scripts (https://github.com/ucdavis-bioinformatics/proc10xG) were used to remove the barcode sequences from the raw reads. SGA-preqc was employed for genome size estimation of *T. cordifolia* using k-mer count distribution method **(Supplementary text S2)** [29]. The de novo assembly was generated by Supernova assembler v.2.1.1 (with maxreads=all options and other default settings) using 499.36 million raw reads [30]. The ‘pseudohap2’ style in Supernova mkoutput was implemented to assemble haplotype-phased genome.

The barcodes of linked reads were processed using Longranger basic v2.2.2 (https://support.10xgenomics.com/genome-exome/software/pipelines/latest/installation) and these processed reads were used by Tigmint v1.2.1 to rectify the mis-assemblies present in Supernova assembled genome [31]. AGOUTI v0.3.3 with quality-filtered transcriptome reads was used to accomplish the initial scaffolding [32]. In order to construct a more contiguous assembly ARCS v1.1.1 with its default parameters was used to provide additional scaffolding and enhance the contiguity of the genome assembly [33]. Using a bloom filter-based method and k-mer value ranging from 30 to 120 with 10 bp interval, Sealer v2.1.5 used the linked reads (barcode processed) for gap-closing in the assembly [34]. Performing scaffolding multiple times could give rise to local mis-assemblies, small indels or distinct base errors which were overwhelmed using Pilon v1.23 that utilized the linked reads (barcode-processed) to increase the assembly quality [35]. The completeness of genome assembly was evaluated with BUSCO v4.1.4 which used embryophyte_odb10 database for the assessment [36]. The additional information about the post-processing of genome assembly is provided in **Supplementary Text S2**.

### Transcriptome assembly

The de novo transcriptome assembly was carried out using RNA-Seq data generated in this study. Trimmomatic v.0.38 was used for processing of raw data reads i.e., adapter removal and quality-filtration **[Supplementary Text S2]** [37]. The de novo transcriptome assembly was constructed using Trinity v2.9.1 with strand-specific option and other default parameters using the processed paired-end reads [38]. A Perl script offered in Trinity software package was utilized to evaluate the assembly statistics.

### Genome annotation

The genome annotation was achieved on the polished assembly (length ≥1,000 bp). This genome was used by RepeatModeler v2.0.1 to construct a *de novo* repeat library [39]. The clustering of obtained repeat sequences was performed using CD-HIT-EST v4.8.1 with sequence identity as 90% and 8 bp seed size [40]. Using the repeat library, RepeatMasker v4.1.0 (RepeatMasker Open-4.0, http://www.repeatmasker.org) was used to soft-mask the genome that was used for the construction of gene set. MAKER pipeline that employs *ab initio-*based gene prediction programs as well as evidence-based approaches for prediction of final gene model was used for genome annotation [41]. As an empirical evidence in MAKER pipeline the de novo transcriptome assembly of *T. cordifolia* and protein sequences of its phylogenetically closer species *Beta vulgaris* (belonging to plant order Caryophyllales) were used. The *ab initio* gene prediction, evidence-based alignments and polishing of alignments were achieved using AUGUSTUS v3.2.3, BLAST and Exonerate v2.2.0, respectively with the MAKER pipeline [42, 43]. The completeness of the coding gene set was also assessed using BUSCO v4.1.4 embryophyte_odb10 database. The tandem repeats detection, de novo tRNAs prediction, de novo rRNAs prediction and miRNAs identification (homology based) were performed using Tandem Repeat Finder (TRF) v4.09, tRNAscan-SE v2.0.7, Barrnap v0.9 (https://github.com/tseemann/barrnap) and miRBase database, respectively [44-46]. The detailed information about genome annotation is provided in **Supplementary Text S3**.

### Orthogroups identification

Among all the eudicot species accessible on Ensembl Plants release 48, a total of 35 species were selected by choosing one species from each offered genus. The 35 eudicot species along with an outgroup species, *Zea mays* were used for the identification of orthologs [47]. The 36 selected species along with the outgroup species were - *Actinidia chinensis, Arabidopsis thaliana, Arabis alpina, Beta vulgaris, Brassica napus, Camelina sativa, Cannabis sativa* female, *Capsicum annuum, Citrullus lanatus, Citrus clementina, Coffea canephora, Corchorus capsularis, Cucumis melo, Cynara cardunculus, Daucus carota, Glycine max, Gossypium raimondii, Helianthus annuus, Ipomoea triloba, Lupinus angustifolius, Malus domestica Golden, Manihot esculenta, Medicago truncatula, Nicotiana attenuata, Olea europaea var. sylvestris, Phaseolus vulgaris, Pistacia vera, Populus trichocarpa, Prunus avium, Rosa chinensis, Solanum tuberosum, Theobroma cacao* Matina 1-6, *Trifolium pratense, Vigna angularis, Vitis vinifera*, and *Zea mays*. The orthogroups were formed using the proteome files of 35 selected eudicot species along with *Zea mays* and MAKER retrieved protein sequences of *T. cordifolia*. Among all the protein sequences, the longest isoforms were retrieved for all the species and provided to OrthoFinder v2.4.1 for orthogroups construction [48].

### Orthologous gene set construction

The orthogroups comprising genes from all 37 species were retrieved from all the identified orthogroups. KinFin v1.0 was used to increase the genes in one-to-one orthogroups that identified and extracted fuzzy one-to-one orthogroups among these retrieved orthogroups [49]. In cases where multiple genes were present for a single species in any orthogroup, the longest gene among them was selected as representative.

### Phylogenetic tree construction

MAFFT v7.480 was used to discretely align all the identified fuzzy one-to-one orthogroups for construction of the phylogenetic tree [50]. The multiple sequence alignments were trimmed to eradicate empty sites, and the alignments were concatenated using BeforePhylo v0.9.0 (https://github.com/qiyunzhu/BeforePhylo). The concatenated alignments were used by RAxML v8.2.12, based on rapid hill climbing algorithm, to create the maximum likelihood-based phylogenetic tree (100 bootstrap values and amino acid substitution model ‘PROTGAMMAAUTO’) [51].

## RESULTS

### Sampling and Sequencing of *T. cordifolia* genome and transcriptome

The *T. cordifolia* plant was brought from a plant nursery in Bhopal, Madhya Pradesh, India and the DNA and RNA extracted from leaf was used for sequencing. Two marker genes: ITS and MatK were amplified and sequenced at in-house sanger sequencing facility for the species identification (**Supplementary Text S1)**. The sequenced reads of these marker genes were aligned using BLASTN that showed the highest identity (98.32% for ITS and 99.41% for MatK) with sequences of *T. cordifolia* available at NCBI database and confirmed the plant species as *T. cordifolia*. We generated 79.4 Gbp of genomic data and 34.7 Gbp of transcriptomic data for the genomic assembly and analysis of *T. cordifolia* **(Table 1 and Supplementary Tables S1-S2)**. The barcode sequences were removed from the raw reads and high-quality reads were selected for further analysis. The de novo assembly generated by Supernova assembler v.2.1.1 using 499.36 million raw reads [30], and the ‘pseudohap2’ style in Supernova mkoutput was implemented to assemble haplotype-phased genome. Since the genome size was not known for this plant, we performed the first genome size estimation for *T. cordifolia* using SGA-preqc processed linked reads, and the genome size was estimated to be 1.13 Gbp. Considering this genome size, the sequenced genomic data amounts to 70.2x genome coverage. After scaffolding, mis-assemblies rectification, gap-filling and polishing *T. cordifolia* genome assembly resulted in 1.01 Gbp assembly size (≥2,000 bp) as the final draft. The %GC of the assembled genome was 35.12%, and a total of 56,342 scaffolds were obtained having the N50 of 50.2 Kbp **(Table 1 and Supplementary Table S3)**. The BUSCO completeness was 78.9% in the final polished *T. cordifolia* genome assembly **(Supplementary Table S4)**. The *de novo* transcriptome assembly was constructed using Trinity v2.9.1 with strand-specific option and other default parameters using the processed paired end reads. The Trinity software assembled 2,764,154 bp of *de novo* transcriptome assembly, and a total of 8,208 transcripts were predicted in the transcriptome assembly.

**Table 1.**
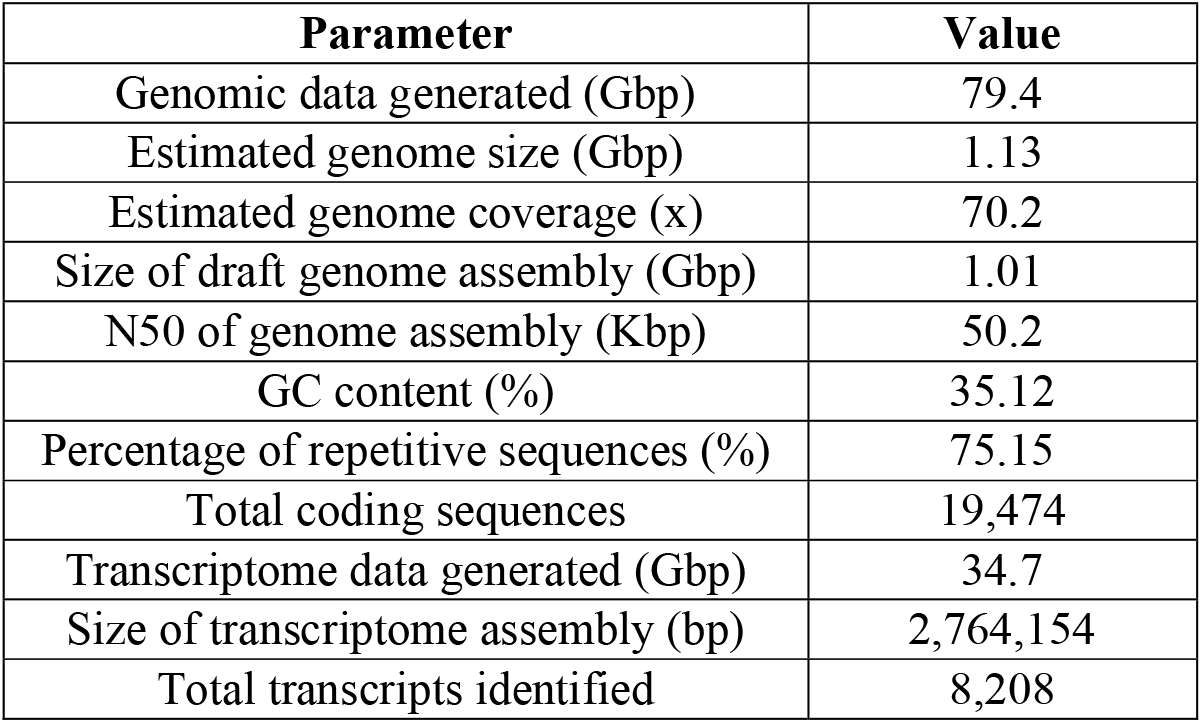
Summary of genome and transcriptome assemblies

### Annotation of genome and gene set formation

The final polished genome assembly was analyzed by RepeatModeler v2.0.1 to construct a *de novo* repeat library consisting of 1,918 repeat families that were further clustered into 1,584 representative repeat family sequences. The repetitive sequences in *T. cordifolia* genome were predicted using RepeatMasker v4.1.0 and obtained 75.15% of repetitive sequences in this genome. Among these repetitive sequences, 73.75% were characterized as interspersed repeats comprising 32.12% retroelements (29.15% of LTR repeats), 2.48% DNA transposons and 39.14% unclassified repeats. The LTR repeats consisted of 25.38% Gypsy/DIRS1 and 3.22% Ty1/Copia elements. About 3.87% of the genome comprised of simple repeats found using TRF v4.09. The interspersed and simple repetitive sequences collectively comprised ∼78% of the total genome. 392 hairpin miRNAs, 1,344 rRNAs and 2,186 tRNAs (decoding standard amino acids) were also predicted among the non-coding RNAs.

The soft-masked genome (generated using RepeatMasker) was used for the construction of gene set using the MAKER pipeline that employs *ab initio-*based gene prediction programs and evidence-based approaches for prediction of final gene model [41]. The de novo transcriptome assembly of *T. cordifolia* and protein sequences of its phylogenetically closer species *Beta vulgaris* (belonging to plant order Caryophyllales) were used as an empirical evidence in MAKER pipeline. The *ab initio* gene prediction, evidence-based alignments, and alignments polishing were achieved using AUGUSTUS v3.2.3, BLAST and Exonerate v2.2.0, respectively in the MAKER pipeline that predicted 19,730 coding sequences in the genome assembly [42, 43]. These coding genes were filtered based on length ≥150 bp and resulted in 19,474 coding gene sequences. The completeness of the coding gene set was assessed using BUSCO v4.1.4 embryophyte_odb10 database on the final gene set that showed 70% of the complete and fragmented BUSCOs to be present in the coding gene set.

### Phylogenetic analysis of *T. cordifolia*

A total of 35 species were selected from the eudicot species accessible on Ensembl Plants release 48 by choosing one species from each offered genus, and included *Actinidia chinensis, Arabidopsis thaliana, Arabis alpina, Beta vulgaris, Brassica napus, Camelina sativa, Cannabis sativa* female, *Capsicum annuum, Citrullus lanatus, Citrus clementina, Coffea canephora, Corchorus capsularis, Cucumis melo, Cynara cardunculus, Daucus carota, Glycine max, Gossypium raimondii, Helianthus annuus, Ipomoea triloba, Lupinus angustifolius, Malus domestica Golden, Manihot esculenta, Medicago truncatula, Nicotiana attenuata, Olea europaea var. sylvestris, Phaseolus vulgaris, Pistacia vera, Populus trichocarpa, Prunus avium, Rosa chinensis, Solanum tuberosum, Theobroma cacao* Matina 1-6, *Trifolium pratense, Vigna angularis*, and *Vitis vinifera*. Thus, the 36 selected species along with one outgroup species (*Zea mays)*, and including *T. cordifolia* were used to detect 162,809 orthogroups using the protein sequences retrieved from 37 species.

The phylogenetic tree (maximum likelihood-based) was constructed using 454 fuzzy one-to-one orthogroups predicted by KinFin v1.0 across the 37 species. The missing or unknown values were predicted by aligning, concatenating and filtering all the fuzzy one-to-one orthogroups. The alignment data was filtered resulting in 422,034 alignment positions. The phylogenetic tree was constructed using this alignment data (filtered) of *T. cordifolia* along with the 35 dicot species available on Ensembl Plants release 48 and an outgroup species, *Zea mays* **(Figure 2)**. The resultant phylogenetic position of *T. cordifolia* was observed as a separate clade from all the other eudicots and monocot outgroup plausibly because the clade to which *T. cordifolia* belongs is considered as basal eudicots that diverged very early from the other eudicots [52, 53].

## DISCUSSION

*T. cordifolia* is known to produce several phytochemicals as secondary metabolites such as alkaloids, terpenoids, tannins, phenolic compounds, steroids, flavonoids, phytosterols, volatile oils, etc. that are responsible for its diverse medicinal properties like anti-oxidant, immunomodulatory, anti-microbial, anti-viral, anticancer, anti-pyretic, anti-inflammatory, hepatoprotective, neuroprotective, anti-osteoporotic, anti-toxic, anti-diabetic, anti-arthritis, anti-ulcer, etc. [18, 54] **(Figure 1)**. Tinosporaside, tembetarine and phenolic compounds were suggested to responsible for its anti-inflammatory, anti-diabetic and anti-oxidant activities, respectively [55-57]. Likewise, a few other compounds from *Tinospora cordifolia* were found accountable for its various activities [58-62]. Being a medicinally important herb with therapeutic applications in multiple health conditions, the genome and transcriptome sequencing of *Tinospora cordifolia* was much needed to gain insights into its genetic information, transcriptome, and functional analysis of pathways associated with the secondary metabolites production responsible for its medicinal properties. This study reported the first draft genome assembly of *T. cordifolia* which is also the first genome from the genus *Tinospora* and its family. This study also deliberates the genome assembly, transcriptome assembly, gene set and phylogeny of *T. cordifolia* that will help in understanding the medicinal properties of this multipurpose plant. The linked read sequencing has recently emerged as a promising technology to significantly increase the contiguity in the genome assembly by increasing scaffold N50 and decreasing the number of scaffolds compared to short read technology, and was successfully used for the sequencing of *T. cordifolia* [63].

**Figure 1.**
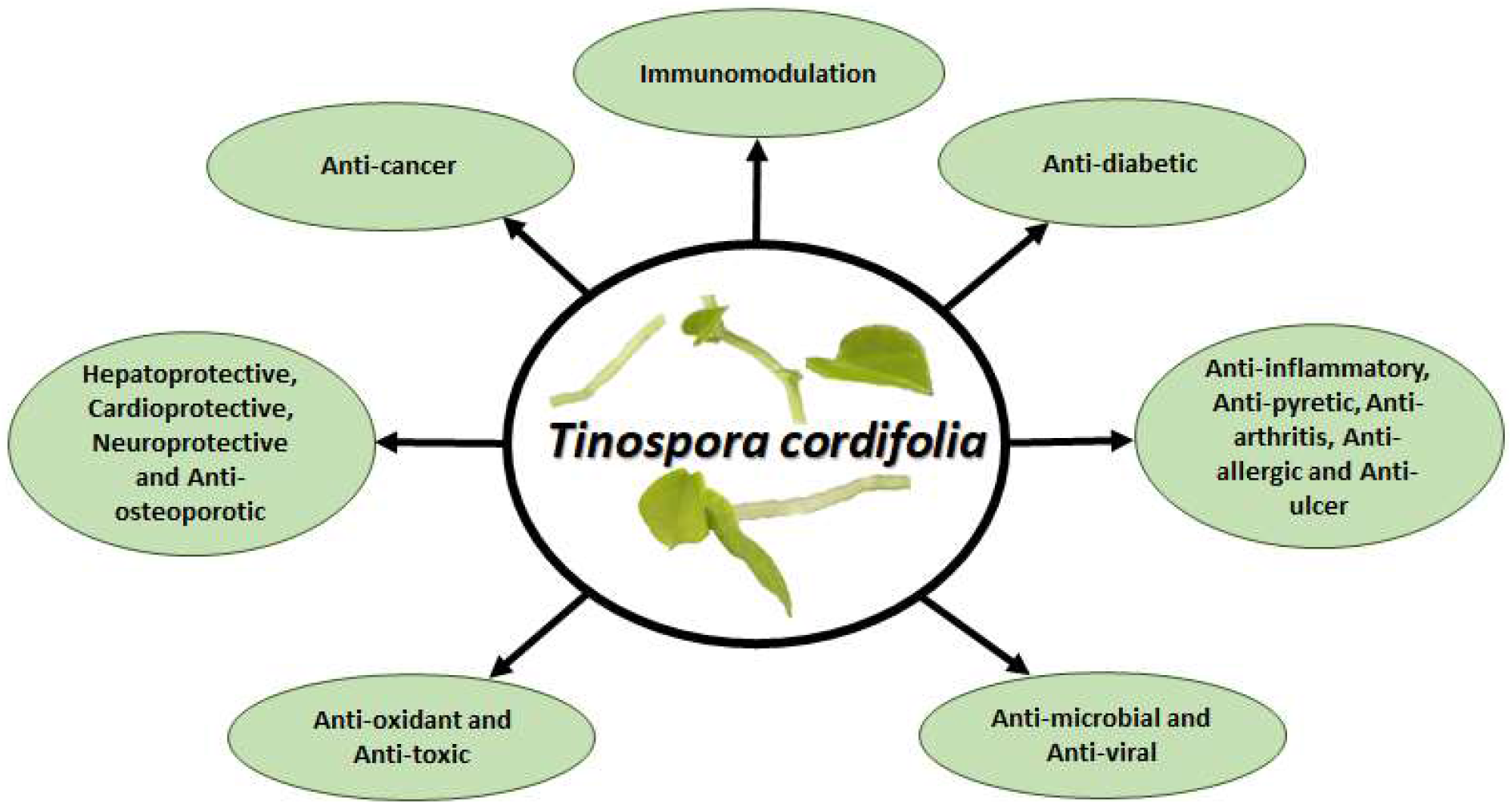
Medicinal properties of *Tinospora cordifolia*

*T. cordifolia* belongs to order Ranunculales and our phylogenetic tree finds its position as a distinct branch separate from all the other eudicots. This could be due to the early-divergence of order Ranunculales from all the other core eudicots. Order Ranunculales is considered as an early-diverging eudicot order and is among a few other eudicot orders (collectively known as basal eudicots) that are found to be sister lineage to the core eudicots [52, 64, 65], which was also observed in the case of *T. cordifolia* that showed early divergence from all other dicot species (**Figure 2**). Thus, the early divergence could be the reason for its distinct position relative to the other eudicots and monocot species. The correctness of this phylogeny is supported by the fact that the species belonging to the same clade shared common nodes, e.g., *Arabidopsis thaliana, Camelina sativa, Brassica napus*, and *Arabis alpina* shared the same clade because they belong to the same order Brassicales, and similarly *Gossypium raimondii, Theobroma cacao* Matina 1-6 and *Corchorus capsularis* belong to order Malvales and shared the same node in the phylogeny. Likewise, the members of order Fabales and order Solanales also shared their respective nodes.

**Figure 2.**
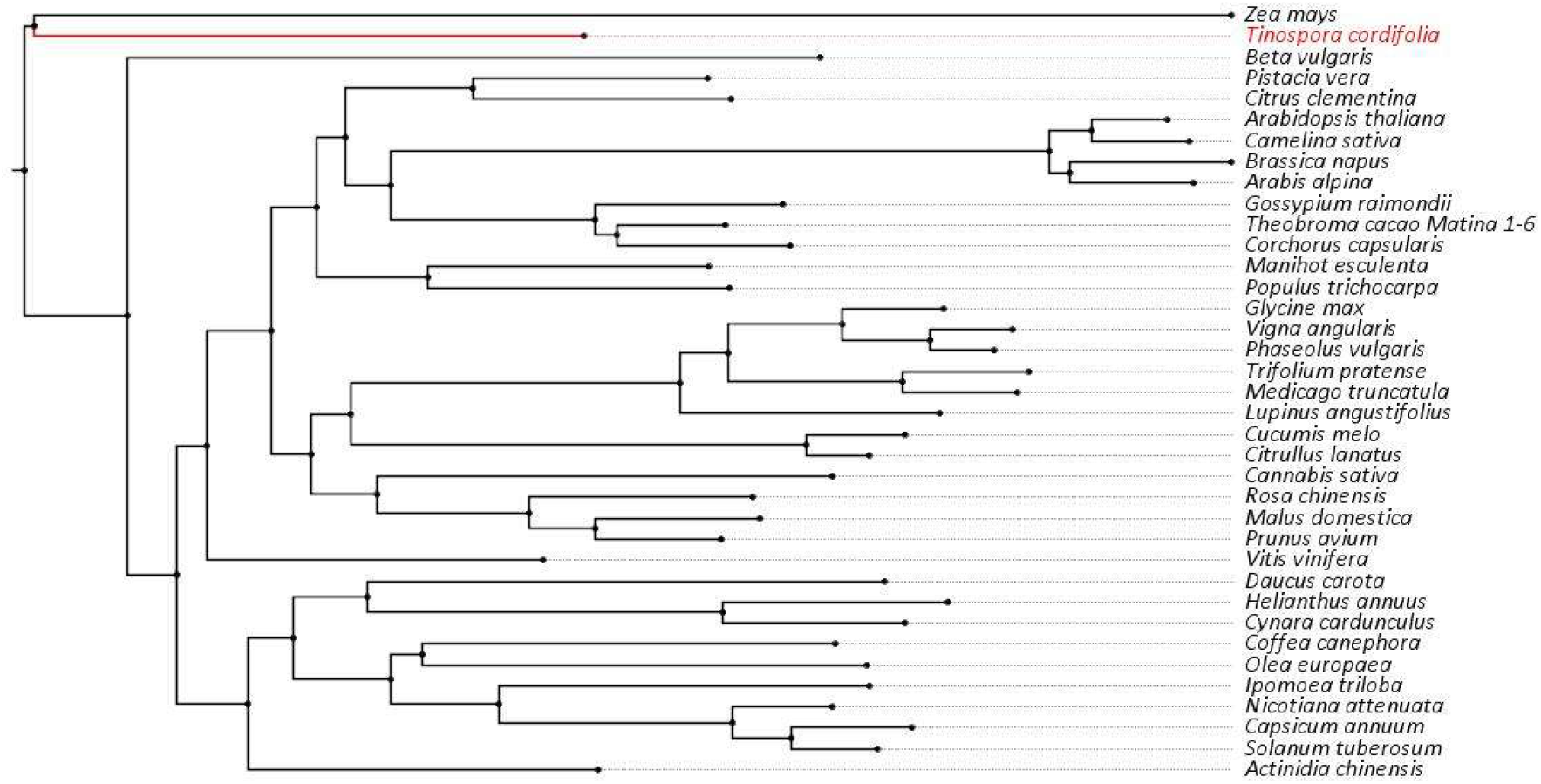
Phylogenetic tree describing the phylogenetic position of *Tinospora cordifolia* with other species

## CONCLUSION

Thus, as apparent from the above literature-based evidences on the medicinal and pharmaceutical properties of this plant, the availability of *T. cordifolia* genome will help in bridging the missing link between its genomic and medicinal properties and provide leads for exploring the genomic basis of these properties. It will also aid in various comparative genomic studies and will act as a reference for the future species sequenced from its genus and family. It will also help in the genome-wide phylogenetic assessments as well as evolutionary analyses on this species. The knowledge of mechanisms and pathways involved in production of its numerous medicinally important secondary metabolites will help in better exploitation of these pathways and resultant metabolites for medicinal purposes and therapeutic applications.

## LIST OF ABBREVIATIONS

COVID: Corona Virus Disease
Gbp: Giga base pair
RNA: Ribonucleic Acid
Mbp: Mega base pair
DNA: Deoxyribonucleic acid
CTAB: Cetyl trimethyl ammonium bromide
tRNA: Transfer RNA
rRNA: Ribosomal RNA
miRNA: micro-RNA
TRF: Tandem repeat finder
ITS: Internal transcribed spacer
MatK: Maturase K
BLAST: Basic Local Alignment Search Tool
BLASTN: Nucleotide BLAST
NCBI: National Center for Biotechnology Information
bp: Base pair
kbp: Kilo base pair
LTR: Long terminal repeats
DIRS1: Dictyostelium intermediate repeat sequence 1

## AUTHORS’ CONTRIBUTIONS

VKS perceived and coordinated the project and collected the plant sample. SM extracted the nucleic acids (DNA and RNA) from the collected sample, prepared the DNA and RNA samples for sequencing, and carried out the species identification assay. AC and VKS conceived the computational outline of the study. AC accomplished all the computational analysis presented in the study. SM, AC and VKS interpreted the results. SM and AC constructed the figures. SM, AC, TS and VKS wrote the manuscript. All the authors have read and approved the final version of the manuscript.

## ACKNOWLEDGEMENT

SM and AC acknowledge Council of Scientific and Industrial Research (CSIR) for supporting research fellowships. TS thanks the Department of Science and Technology and IISc Bangalore for providing the KVPY scholarship. The authors also thank IISER Bhopal for providing the intramural research funds.

## DATA AVAILABILITY

The raw data has been submitted on NCBI database with BioProject accession number PRJNA749156, Biosample accession SAMN20355817, and SRA accessions SRR15221491 and SRR15221490.

