## Supplementary Information for "Genome sequencing and assembly of *Tinospora cordifolia* (Giloy) plant"

### SUPPLEMENTARY TABLES

**Supplementary Table S1. Detailed information of genomic linked-read data for *Tinospora cordifolia***

| Average Read Length (bp) | Total Number of Reads (Millions) | Total Generated Data (Gb) | Coverage |
|---|---|---|---|
| 159 | 249,680,248(x2) | 79.4 | ~70x |

The sequencing coverage was calculated considering 1.13 Gbp as the estimated genome size

**Supplementary Table S2. Description of *Tinospora cordifolia* transcriptome data**

| Average Read Length R1 (bp) | Average Read Length R2 (bp) | Total Number of Read pairs | Total Number of Bases in R1 (bp) | Total Number of Bases in R2 (bp) | Total Number of Bases (bp) |
|---|---|---|---|---|---|
| 161 | 161 | 107,858,957 (x2) | 17,365,292,077 | 17,365,292,077 | 34,730,584,154 |

**Supplementary Table S3. Statistical details of final *de novo* assembled draft genome using 10x genomics reads of *Tinospora cordifolia***

| Parameter | Value |
|---|---|
| Number of contigs (>= 1,000 bp) | 76,178 |
| Number of contigs (>= 2,000 bp) | 56,342 |
| Number of contigs (>= 5,000 bp) | 34,423 |
| Number of contigs (>= 10,000 bp) | 20,212 |
| Number of contigs (>= 25,000 bp) | 10,599 |
| Number of contigs (>= 50,000 bp) | 5,569 |
| Total length (>= 1,000 bp) | 1,040,091,788 |
| Total length (>= 2,000 bp) | 1,012,254,697 |
| Total length (>= 5,000 bp) | 940,788,363 |
| Total length (>= 10,000 bp) | 840,582,365 |
| Total length (>= 25,000 bp) | 691,969,894 |
| Total length (>= 50,000 bp) | 508,118,085 |
| Largest contig | 786,413 |
| GC (%)(>= 2,000 bp) | 35.12 |
| N50 (>= 2,000 bp) | 50,214 |
| L50 (>= 2,000 bp) | 5,530 |

| | |
|---|---|
| Number of N's per 100 kbp (>= 2,000 bp) | 1106.21 |

**Supplementary Table S4. BUSCO measurements of *Tinospora cordifolia* genome**

| Parameters | Supernova v2.1.1 assembled raw genome | Final draft genome (≥1,000 bp) |
|---|---|---|
| Complete BUSCOs (C) | 1,256 (77.8%) | 1,274 (78.9%) |
| Fragmented BUSCOs (F) | 181 (11.2%) | 169 (10.5%) |
| Missing BUSCOs (M) | 177 (11%) | 171 (10.6%) |
| Total BUSCO groups searched | 1,614 | 1,614 |

The embryophyta_odb10 database was used as the reference database for BUSCO analysis.

### SUPPLEMENTARY TEXTS

#### Supplementary Text S1.

#### Sample collection and Nucleic acids Extraction

The plant was brought from a nursery situated in Bhopal, Madhya Pradesh, India (23.2599° N, 77.4126° E). The leaves were collected from the plant and were used for extraction of nucleic acids (DNA and RNA). The leaves were cleaned and homogenised in liquid nitrogen using pre-cooled autoclaved mortar-pestle. For DNA extraction, the homogenized leaves were washed once with 70% ethanol and twice with autoclaved distilled water in order to remove any contaminants which can hinder the extraction process. This treated powdered leaves were lysed using Carlson lysis buffer [100mM Tris HCl, 2% Cetyl Trimethyl Ammonium Bromide (CTAB), 1% PEG 8000, 1.4M NaCl and 20 mM EDTA (pH 9.5)] [1]. After adding β-mercaptoethanol (2.5 µL) to each ml of lysis buffer it was pre-heated for 30 mins at 65˚C. This pre-heated buffer supplemented with 2 µL and 25 µL of RNase A (20 mg/mL) and Proteinase K (20 µL/mL), respectively was added to the homogenized leaves and incubated for 1 hr at 65˚C. The tubes were intermediately mixed by inverting them with the aim of longer DNA fragments. At all the steps, the tubes were mixed only by inversion and centrifuged at 4˚C. The tubes were cooled to room temperature followed by addition of chloroform (1 mL) to the collected supernatant and centrifuged for 15 mins. The aqueous layer was collected in a new centrifuge tube and added 0.7x volume of isopropanol (ice-cold) for DNA precipitation. The tubes were mixed and kept at -20˚C for overnight in order to facilitate the precipitation. Next day the tubes were centrifuged for 45 mins to pellet down the DNA. The pelleted DNA was washed with 70% ethanol by centrifuging for 30 mins and supernatant was removed followed by air drying of pellet to remove all the residual ethanol. The DNA pellet was dissolved in 100 µL of Nuclease free water. The quality of extracted DNA was evaluated on NanoDrop™8000 Spectrophotometer (ThermoFisherScientific, USA) as well as 1% agarose gel electrophoresis. The DNA quantification was carried out on Qubit 2.0 fluorometer using Qubit dsDNA BR assay kit (Invitrogen, USA).

For RNA extraction, 1 mL of TRIzol reagent (Invitrogen, USA) was added to 100 mg of homogenized leaves and mixed for 5 mins using a vortexer. After addition of chloroform (200 µL), the tubes were briefly vortexed and incubated for 10 mins at room temperature. All centrifugation steps were carried out at a speed of 12,000xg and 4˚C temperature. A centrifugation for 15 mins was provided to separate the aqueous layer. The separated aqueous layer was treated with ice-cold isopropanol (500 µL) and incubated for 10 mins at room temperature. A centrifugation for 10 mins was given in order to pellet down the RNA. The RNA pellet was washed with 1 mL of 70% ethanol and the residual ethanol was removed by incubating for 30 mins at 37 ˚C. Nuclease free water (50 µL) was used for resuspending the RNA pellet followed by an incubation of 10 mins at 55˚C [2]. The quality and quantity of extracted RNA were evaluated on NanoDrop™8000 Spectrophotometer (ThermoFisherScientific, USA) and 1% agarose gel electrophoresis, and Qubit 2.0 fluorometer with Qubit ssRNA HS assay kit (Invitrogen, USA), respectively.

**Species Identification**

Two genes: one nuclear (Internal Transcribed Spacer ITS) and other plastid gene (Maturase K MatK) were used to identify the species of the collected sample. The ITS amplification was done using forward primer 5′-TCCGTAGGTGAACCTGCGG-3′ and reverse primer 5′-TCCTCCGCTTATTGATATGC-3′. Similarly, the forward and reverse primers used for MatK gene amplification were 5’-CGATCTATTCATTCAATATTTC-3’ and 5’-TCTAGCACACGAAAGTCGAAGT-3’, respectively. The PCR programme used were as follows:

1. For ITS gene amplification:
   Initial denaturation: 94 ˚C for 3 mins
   35 cycles of –
   (i) Denaturation at 94 ˚C for 1 min
   (ii) Annealing at 55 ˚C for 1 min
   (iii) Extension at 72 ˚C for 2.5 mins
   Final extension: 72 ˚C for 10 mins

2. For MatK gene amplification:
   Initial denaturation: 95 ˚C for 3 mins
   35 cycles of –
   (iv) Denaturation at 95 ˚C for 30 sec
   (v) Annealing at 50 ˚C for 3 mins
   (vi) Extension at 72 ˚C for 1.15 mins
   Final extension: 72 ˚C for 7 mins

Taq Polymerase (Invitrogen, USA) and Paq5000 polymerase (Agilent, Santa Clara, USA) was used employed for MatK and ITS amplification, respectively. The amplification was checked by running the amplicons using 2X agarose gel electrophoresis. The amplicons were purified and sequenced on a sanger sequencer at the in-house facility. The sequences were checked for alignment using BLASTN which showed highest identity with *Tinospora cordifolia* that confirmed the species.

**DNA and RNA Sequencing**

The library for genome sequencing was prepared on Chromium controller (10x Genomics, CA, USA) using the extracted DNA. The DNA fragments were barcoded on this instrument using Gel bead kit v2 and Chromium Genome Library kit (10x Genomics, CA, USA) as per the manufacturer’s instructions. The library for transcriptome sequencing was prepared using TruSeq Stranded Total RNA Library Preparation kit with Ribo-Zero Plant workflow (Illumina Inc., CA, USA). The quality and quantity of both the libraries were assessed on Tapestation 4150 (Agilent,

Santa Clara, CA) using high sensitive D1000 screentapes and Qubit 4.0 fluorometer using Qubit ds DNA HS assay kit (Invitrogen, USA), respectively. Finally, paired end reads were generated by sequencing both the libraries on Novaseq 6000 (Illumina, Inc., United States).

**Supplementary Text S2.**

**Genome size estimation**

An array of python scripts (https://github.com/ucdavis-bioinformatics/proc10xG) was utilized to cut the barcode sequences from raw linked reads. The process_10xReads.py script was used on all linked reads with its default parameters to extract the barcodes. Based on the barcode status of reads filter_10xReads.py script was used to filter the reads.

SGA-preqc employed a method based on k-mer count distribution for genome size estimation. It removed the low occurring error susceptible k-mers [3]. Initially, filtered linked reads were pre-processed by SGA in paired-end mode followed by indexing of pre-processed reads using 'ropebwt' indexing algorithm with '--no-reverse' option. Lastly, SGA preqc was executed with its default settings for the genome size estimation of *Tinospora cordifolia*.

**Assembling and polishing of genome**

Supernova v2.1.1 used 499.36 million linked reads (not pre-processed) which is equivalent to 70x sequencing coverage for the *de novo* genome assembly. Supernova v2.1.1 generated haplotype-phased fasta assembly file by setting maxread=all option and Supernova mkoutput as 'pseudohap' style [4]. The barcodes of linked reads were processed using Longranger basic v2.2.2 (https://support.10xgenomics.com/genome-exome/software/pipelines/latest/installation) and these processed reads were used by Tigmint v1.2.1 to rectify the mis-assemblies present in the genome assembled by Supernova [5]. The indexing of assembled genome was performed that followed by mapping of linked reads (barcode-processed) using bwa-mem and ".bam" file was generated using samtools v1.9. The tigmint-molecule used this ".bam" to create the ".bed" file. The corrected assembly was generated by tigmint-cut using the ".bed" file to remove regions of mis-assembly.

Longranger align v2.2.2 (https://support.10xgenomics.com/genome-exome/software/pipelines/latest/installation) performed the initial scaffolding by mapping the linked reads to the corrected genome assembly. The samtools v1.9 generated ".bam" file along with ARCS v1.1.1 and LINKS v1.8.6 (with default parameters) were used to construct the genome assembly of *Tinospora cordifolia* with increased contiguity [6, 7]. The *de novo* transcriptome assembly was performed by AGOUTI v0.3.3 using filtered RNA-seq reads [8]. The previously scaffolded genome was mapped with RNA-seq reads using bwa-mem and the ".bam" file was generated using samtools v1.9 [9, 10]. AGOUTI generated additionally scaffolded genome assembly using above obtained ".bam" file and ".gff3" file derived from AUGUSTUS v3.2.3 [11]. Using a bloom filter of size 950 GB and k-mer value ranging from 30 to 120 with 10 bp interval, the linked reads (barcode processed) were utilized by Sealer v2.1.5 for gap-closing in the assembly [12]. Using bwa-mem this gap-closed genome assembly was mapped for a second time with barcode-processed linked reads and ".bam" file was generated using samtools v1.9 [9, 10]. The genome assembly was polished by Pilon v1.23 using this ".bam" file to increase the quality of genome assembly [13].

**Pre-processing of transcriptome data**

The pre-processing of transcriptome data was carried out using Trimmomatic v0.38 [14]. Adapters were trimmed allowing two as maximum mismatches in a 16 bp seed match. It also used thresholds as 30 for palindrome clip and 10 as simple clip. The both ends of a read were trimmed by setting the quality threshold value to 15. If the average quality score for a four-base window was found to be less than 15 then sliding window for reads was trimmed. Elimination of all reads with <40 bases was done.

**Supplementary Text S3.**

**Identification of tandem repeats**

Tandem Repeat Finder (TRF) v4.09 was used on the final polished draft genome (≥1,000 bp) of *Tinospora cordifolia* for identification of tandem repeats. The parameters used for this were matching weight =2, match probability = 80%, mismatching penalty =7, minimum alignment score = 50, indel probability = 10%, indel penalty =7, and maximum period size = 2000 [15].

**Transfer RNAs (tRNAs) identification**

The *de novo* prediction of transfer RNAs in the final polished draft genome assembly (≥1,000 bp) was carried out using tRNAscan-SE v2.0.7 with its default parameters [16]. This prediction found 2,269 tRNAs in the *Tinospora cordifolia* genome assembly and classified them as 2,186 tRNAs translating Standard 20 amino acid, 1 possible suppressor tRNA, 7 tRNAs with undetermined isotypes and 75 predicted pseudogenes. Along with these, 24 tRNAs with introns were identified.

**microRNAs identification**

The hairpin miRNAs were identified based on homology using miRBase database [17]. This analysis identified 38,589 hairpin miRNAs which were clustered by CD-HIT-EST v4.8.1 with sequence identity of 90%. 22,365 non-redundant sequences were obtained by clustering [18]. These 22,365 sequences were used to identify the hairpin miRNAs using BLASTN with parameters such as identity and e-value of 80% and 1e-03, respectively [19].
